# Prescription Opioid Distribution After the Legalization of Recreational Marijuana in Colorado, 2007-2017

**DOI:** 10.1101/702811

**Authors:** Amalie K. Kropp, Stephanie D. Nichols, Daniel Y. Chung, Kenneth L. McCall, Brian J. Piper

**Affiliations:** Geisinger Commonwealth School of Medicine, Scranton, PA 18509; University of New England, Portland, ME 04005

**Keywords:** Marijuana, opioids, Colorado, recreational

## Abstract

**Importance:** Opioid related overdoses and overprescribing continue to be an ongoing issue within the United States. Further consideration of nonopioid alternatives as a substitute to treat chronic noncancer pain and in the treatment of opioid use disorders (OUD) is warranted.

**Objective:** To examine the association between the legalization of Colorado’s recreational marijuana and prescription opioid distribution trends. Two states that have not legalized recreational marijuana were selected for comparison.

**Methods:** The United States Drug Enforcement Administration’s Automation of Report and Consolidated Orders System (ARCOS) was used to examine nine pain medications: oxycodone, fentanyl, morphine, hydrocodone, hydromorphone, oxymorphone, tapentadol, codeine, meperidine and two OUD medications: methadone and buprenorphine from 2007-2017 in Colorado, Utah, and Maryland. The drug weights were extracted, examined, and graphed. Medications were converted to their oral morphine milligram equivalents (MME) using standard conversion factors.

**Results:** Colorado reached a peak of pain MME weight in 2012 and had an −11.66% reduction from 2007 to 2017. During the same interval, Utah had a +9.64% increase in pain medication distribution and Maryland, a −6.02% reduction. As for medications used for OUD, Colorado, Utah, and Maryland had +19.42% increase, −31.45% reduction, and +66.56% increase, respectively. Analysis of the interval pre (2007-2009) versus post (2013-2017) marijuana legalization was completed. Statistically significant changes were observed for Colorado (P=0.033) and Maryland (P=0.007), but not Utah (P=0.659) for pain medications. Analysis of the OUD medications identified significant changes for Colorado (P=0.0003) and Maryland (P=0.0001), but not Utah (P=0.0935). Over the decade, Colorado’s opioid distribution was predominantly (72.49%) for pain with one-quarter (27.51%) for an OUD. Utah distributed 61.00% for pain and 39.00% for OUD. However, Maryland was one-third (37.89%) for pain but over-three-fifths (62.11%) for an OUD.

**Conclusion:** There has been a significant decrease in the prescription opioid distribution after the legalization of marijuana in Colorado. This finding was particularly notable for opioids indicated predominantly for analgesia such as hydrocodone, morphine and fentanyl. Colorado had a larger decrease in opioid distribution after 2012 than Utah or Maryland. Therefore, marijuana could be considered as an alternative treatment for chronic pain and reducing use of opioids. Also, when combined with other novel research, it may also reduce the overdose death rate. Additional research with more comparison states is ongoing.

## Introduction

An epidemic is plaguing the United States regarding the misuse of prescription opioids over the last fifteen years. The opioid epidemic stems from the early 1990s when the medical community started to recognize pain as a fifth vital sign.^1^ This philosophical shift caused the implementation of national initiatives to improve pain-related care in 2001. Specifically, the Joint Commission released guidelines that affected physician prescribing behavior on a state and national level. The Drug Enforcement Administration (DEA) found a marked increase in the total number of opioids prescribed every year since the mid-1990s^2^, leading up to the national peak in opioid distribution in 2012. Opioid prescribing increased from 148 million prescriptions in 2005 to over 206 million by the end of 2011.^3^ Although the intent of these guidelines was to improve pain care, there was no significant increase in the quality of pain management.^1^ This has had long-lasting and devasting effects that have rippled throughout the entire country as more people became dependent and overdosed on opioids. While there have been downward trends in prescribing of opioids after 2012, patients are still dying. Ninety Americans die each day due to overdoses^4^. There has been an increase of 320% between 2000 and 2015 in opioid-related mortality and it is still three-times higher than in 1999.^5^ Physicians and other healthcare providers have the responsibility to treat their patient’s non-cancer pain, while also considering nonopioid alternatives.

Since California first legalized medical marijuana in 1996, 33 states and the District of Columbia have passed laws broadly legalizing marijuana, either medically or recreationally, as of June 2019. Washington D.C. and ten other states have expanded to recreational marijuana use.^6^ With the endorsement of the states, more objective, clinical evidence is surfacing that marijuana can be used to manage chronic pain^7^, reduce overdose mortality rates^8,9^, treat opioid withdrawal^10^, and decrease opioid prescribing rates.^5,11,12^

Marijuana has a much lower risk of addiction and virtually no overdose danger, which is in stark contrast to opioids.^13,14,15,16^ In Jan. 2017, the National Academies of Sciences, Engineering and Medicine released a peer-reviewed, comprehensive review showing, “conclusive evidence” that marijuana can be used safely and effectively to treat chronic pain.^17^ According to a Pew research center poll conducted in 2018, 62% of Americans are supportive of legalized marijuana for medical purposes, which has doubled in over a decade from 31% in 2000.^18^ The popularity of marijuana is quickly rising in the 50 and older age group.^19^ This age group may be most likely to experience chronic pain related conditions and are open to the analgesic effects of marijuana.^7^ Overall, marijuana is gaining strong support politically. This rising acceptance also helps dispels myth of marijuana being a gateway drug, decreasing the stigma of this alternative treatment.

To date, there has been little to no research conducted on the effects of adult use marijuana laws on opioid distribution. In November 2000, 54% of Colorado voters approved Amendment 20, implementing the legalization of medical marijuana.^20^ Twelve years later, Colorado approved Amendment 64, legalizing adult-use or recreational marijuana.^20^ By January of 2014, dispensaries were opened to the public. This research compares medical opioid distribution in Colorado with two states that have not legalized recreational marijuana.

## Methods

### Procedures

The Automation of Report and Consolidated Orders System (ARCOS) is a federal program created because of the 1970 Controlled Substances Act that collects data on narcotics in Schedules II to III from hospitals, narcotic treatment programs (NTPs, also known as methadone programs), and pharmacies.^2^ The program is overseen by the Drug Enforcement Administration (DEA) with the goal of preventing, detecting, and investigating controlled pharmaceuticals.^2^ Nine opioid pain medications: oxycodone, fentanyl, morphine, hydrocodone, hydromorphone, oxymorphone, tapentadol, codeine, and meperidine and two primarily opioid use disorder (OUD) medications: methadone and buprenorphine, were examined from 2007-2017 in Colorado, Utah, and Maryland. Colorado and Maryland had similar demographics in terms of population size, percent of house owners, percentage of bachelor’s degrees or higher education holders, and uninsured rates gathered from the US Census 2018 data. Utah was chosen as the most geographically similar state with similar Body Mass Index (BMI) and median household income (Table 1).^21^ These social determinants of health are important to consider when looking at opioid distribution at a state level. Social, regional, and economical factors play substantial parts in opioid use and misuse when examining the multifactorial realm of pain and treatment options. Institutional Review Board approval was provided by the University of New England.

### Statistical Analysis

All eleven medications were converted to their oral morphine milligram equivalents (MME). This enabled the agents to be compared despite their differences in relative potency. Oral MME conversions were completed using the following multipliers: oxycodone 1.5, fentanyl 75, morphine 1, hydrocodone 1, hydromorphone 4, oxymorphone 3, tapentadol 0.4, codeine 0.15, meperidine 0.1, methadone 8, and buprenorphine 10.^22^ Heat maps of three-digit zip codes were prepared using QGIS; other figures were created with GraphPad Prism, version 8.1. Population data was taken from Statista. T-tests compared pre-marijuana legalization (2007-2012) and post-marijuana legalization (2013-2017) for the nine pain medications and two OUD medications. T-tests were calculated with GraphPad QuickCalcs. A p < .05 was considered statistically significant but trends (p < .10) were also noted.

## Results

Figure 1 shows the total MMEs of all eleven opioids, opioids primarily used for OUD and opioids used for pain from 2007 to 2017. Maryland had the highest weight and peaked in 2011 at 12,167 kg MME for all eleven opioids, over twice the weight than the comparison states. Colorado and Utah, at their peaks only reached 5,029 kg MME in 2012 and 3,429 kg in 2015, respectively. Over the decade, Maryland had an increase in methadone and buprenorphine, and about the same amount of distribution of pain medications. Utah remained relatively constant in all categories over the decade. Colorado, similar to Maryland, also experienced an increase distribution of OUD medications, an increase in all opioids, and small elevations in pain medications.

**Figure 1.**
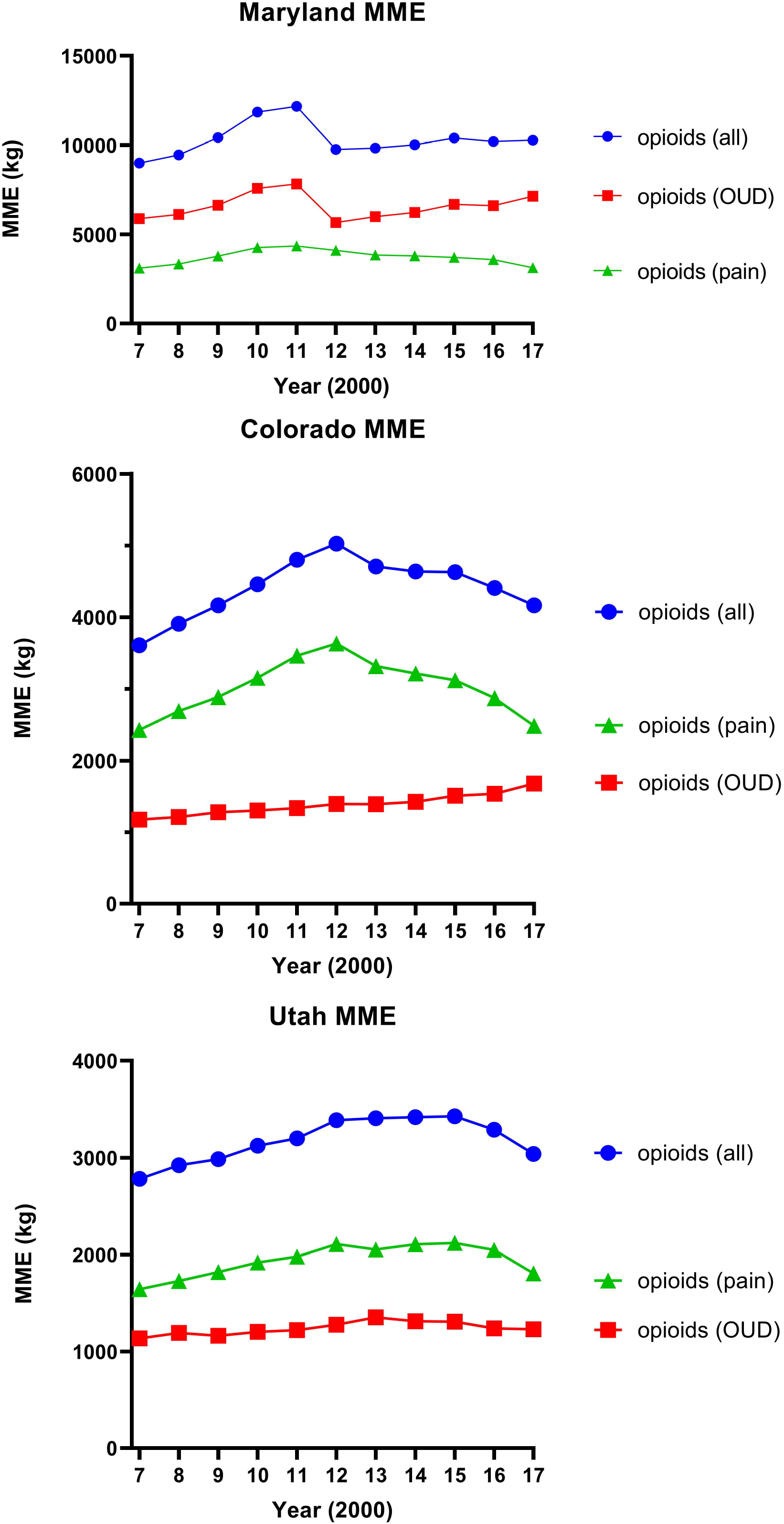
Medications for pain (oxycodone, fentanyl, morphine, hydrocodone, hydromorphone, oxymorphone, tapentadol, codeine, meperidine), Opioid Use Disorder (OUD, methadone and buprenorphine), and all 11 drugs in their morphine milligram equivalents in kilograms per year in Colorado, Maryland and Utah from 2007 to 2017.

Figure 2 shows the MME of each opioid. Oxycodone and methadone were the most distributed in all states. Methadone in Maryland peaked in 2011 at 7,237 kg. In contrast, methadone in Colorado peaked at 1,384 kg in 2017 and, in Utah, 1,054 kg in 2013. Tapentadol, codeine, and meperidine were the least distributed. Buprenorphine increased over the decade in all states.

**Figure 2.**
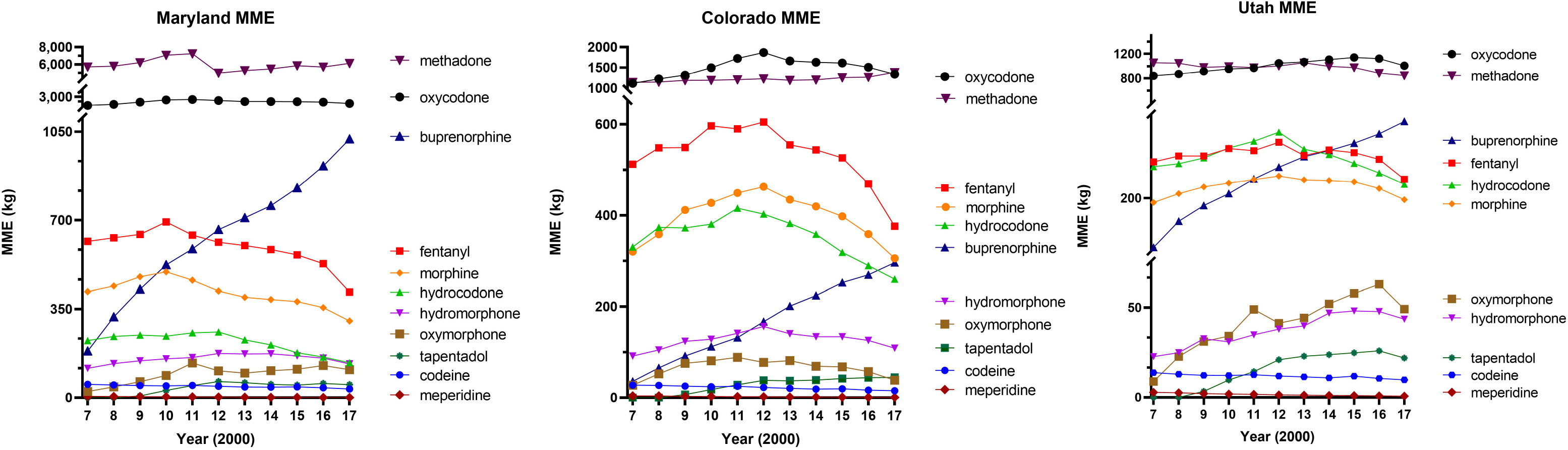
Morphine milligram equivalents of 11 drugs in Colorado, Utah, and Maryland from 2007 to 2017.

Figure 3 shows heat maps of the relative MME per zip code, per population in kg weight in 2017. Colorado’s densest zip code was 802, surrounding Denver, the state capital with over 1.24 metric tons of opioid in MME distributed. Baltimore, Maryland, with the zip code 212, was the densest, with over 4 metric tons of opioids distributed. Interestingly, Utah had two zip codes with similar MME weights, unlike either Colorado or Maryland. Zip codes, 840 and 841, both around Salt Lake, had MME weights of 1.15 metric tons. Based on US Census 2010 data Denver (zip code-802) had a population of 600,158. Baltimore, Maryland (zip code-212) had 620,961 and Salt Lake City, Utah (zip code-841) had a population of 186,440.

**Figure 3.**
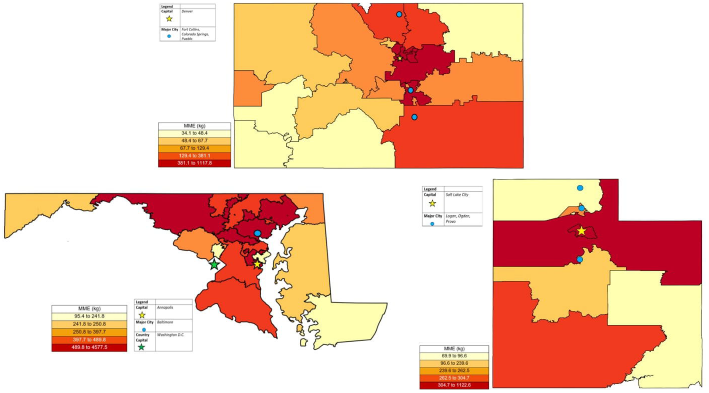
Heat maps showing morphine milligram equivalents per zip code, per population in kilogram weight. Capital and major cities are shown. Population data from Statista. Maps created with QGIS.

The MMEs of the nine pain medications per person were graphed over the decade within each state in Figure 4. Colorado had an −11.66% reduction from 2007 to 2017. Utah had a +9.64% increase and Maryland, a −6.02% reduction in pain medication distribution. The two OUD medications were also looked at. Colorado, Utah, and Maryland had +19.42% increase, −31.45% reduction, and +66.56% increase, respectively. This sizable increase in Maryland was a unique finding, as neither of the other states showed this large of a change. Although, Colorado and Maryland were trending up over the last decade in regard to OUD medication distribution, Utah had a significant decrease in the same timeframe.

**Figure 4.**
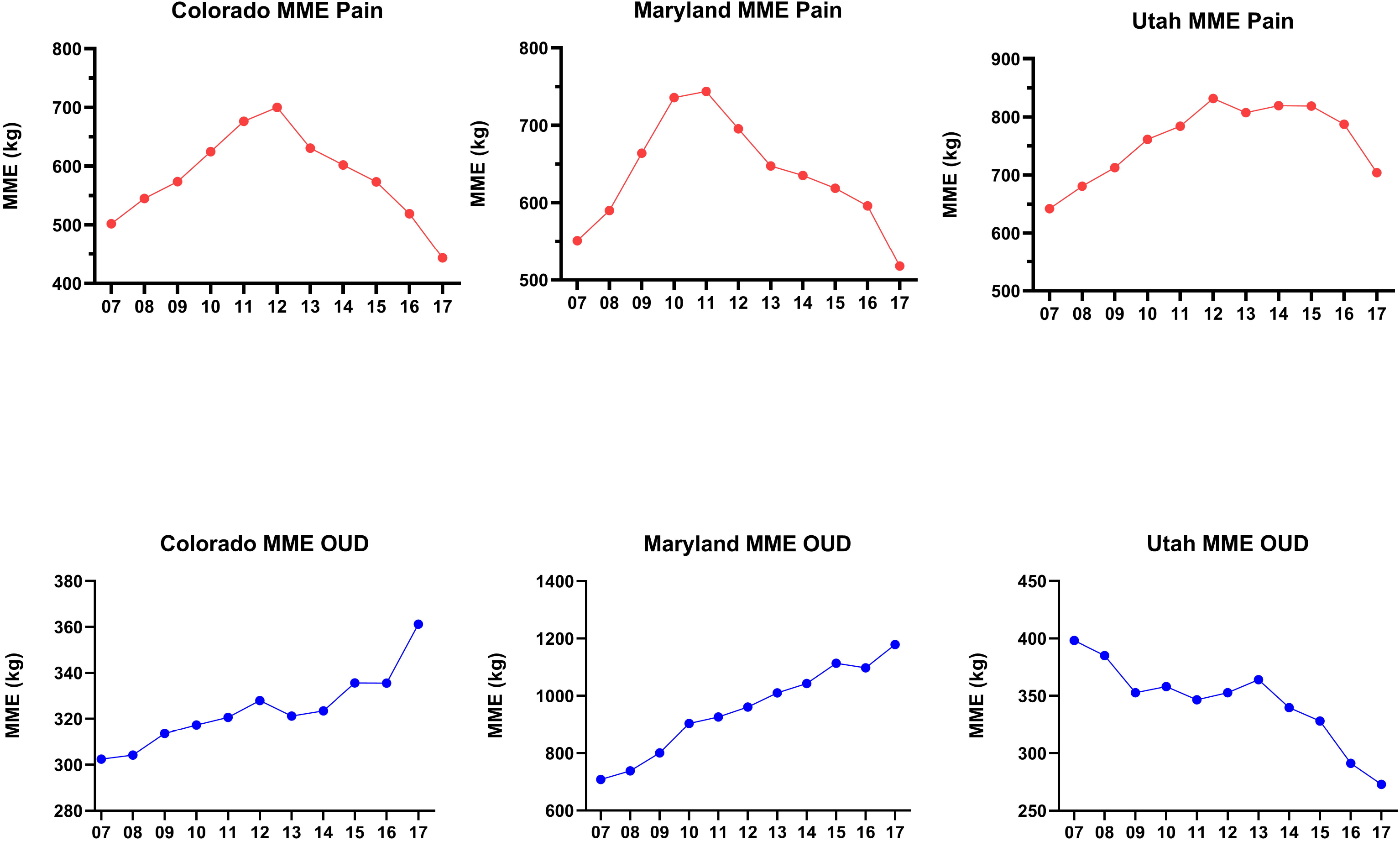
Morphine milligram equivalents of nine pain opioid in Colorado, Utah, and Maryland per person from 2007 to 2017. Colorado had an −11.66% reduction from 2007 to 2017. Utah had a +9.64% increase in pain medication distribution and Maryland, − 6.02% reduction (top). Morphine milligram equivalents of 2 OUD drugs in Colorado, Utah, and Maryland per person from 2007 to 2017. Colorado, Utah, and Maryland had +19.42% increase, −31.45% reduction, and +66.56% increase, respectively (bottom).

Figure 5 highlighted the percent of opioid use for pain and OUD, by their oral morphine milligram equivalents in Colorado, Maryland and Utah from 2007 to 2017. Colorado and Utah showed a similar outcome with 79.49% pain medications and 61.00%, respectively. Maryland ended up reveling an opposite trend. Maryland has only 37.89% pain medication distributed, and 62.11% OUD medications distributed over the decade. Colorado and Utah only had 27.51% and 39.00% of OUD medications dispersed.

**Figure 5.**
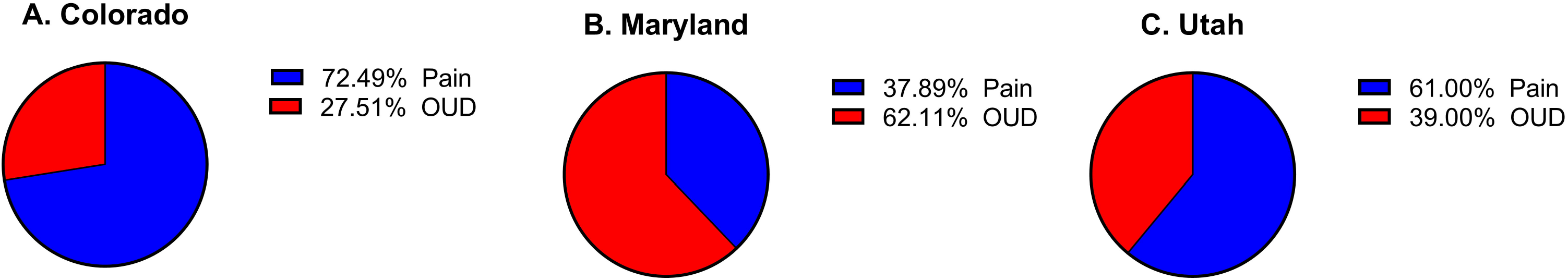
Percent of opioid use for pain (oxycodone, fentanyl, morphine, hydrocodone, hydromorphone, oxymorphone, tapentadol, codeine, meperidine) and Opioid Use Disorder (OUD, methadone and buprenorphine), by their oral morphine milligram equivalents in Colorado, Maryland and Utah from 2007 to 2017.

T-tests compared pre-marijuana legalization (2007-2012) and post-marijuana legalization (2013-2017) for the nine pain medications and two OUD medications in Figure 6. These years are referencing the legalization of recreational marijuana in Colorado. There were statistically significant reductions for Colorado (P=0.033) and Maryland (P=0.007), but not Utah (P=0.659). Analysis of the OUD medications found statistically significant changes for Colorado (P=0.0003) and Maryland (P=0.0001), but not Utah (P=0.0935).

**Figure 6.**
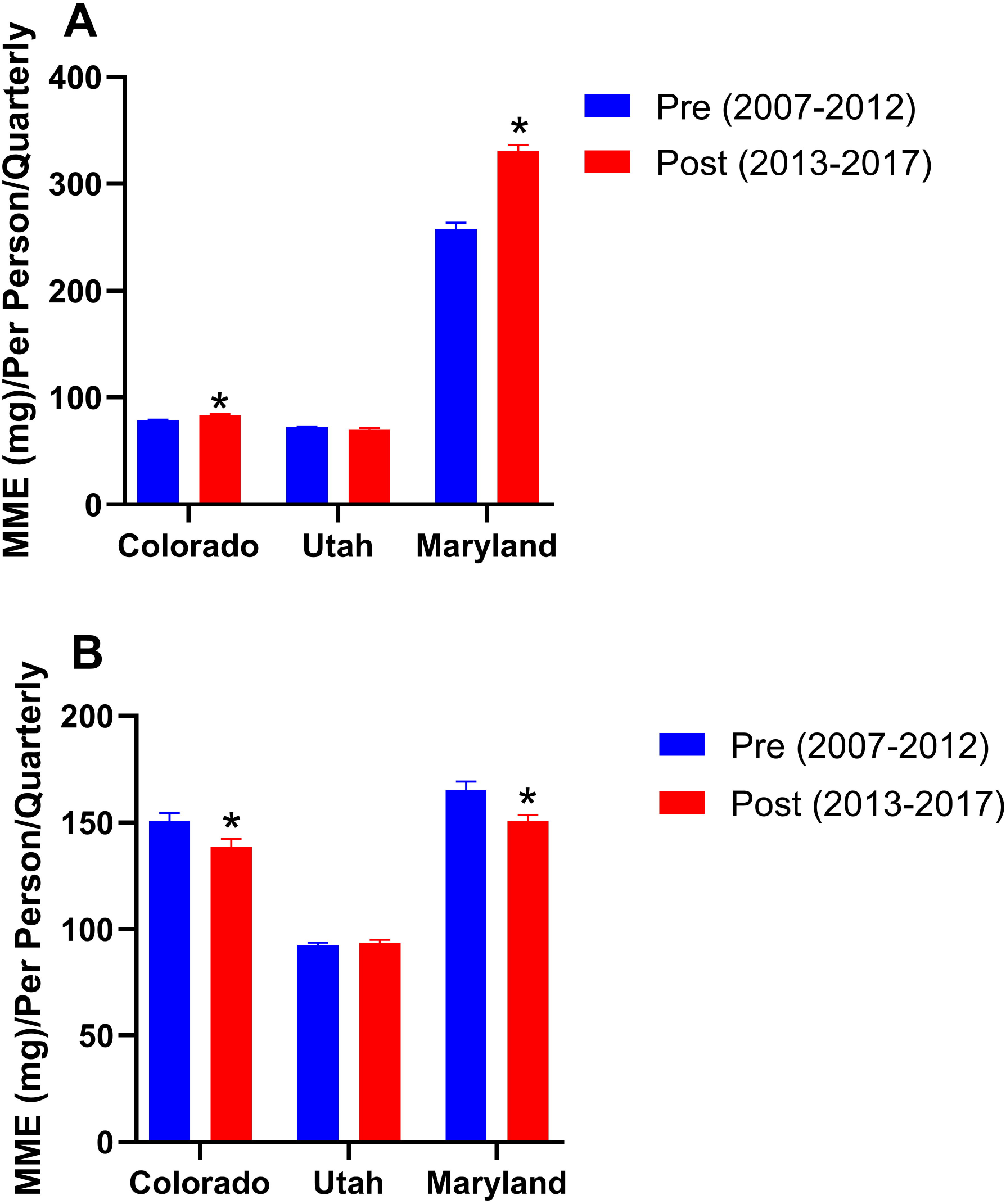
T-test analysis of the nine pain medications found statistically significant reductions for Colorado (P=0.033) and Maryland (P=0.007), but not Utah (P=0.659). T-test analysis of the two Opioid Use Disorder (OUD) medications found statistically significant changes for Colorado (P=0.0003) and Maryland (P=0.0001), but not Utah (P=0.0935).

## Discussion

There are several important findings of this novel pharmacoepidemiology and public policy research that supports recreational marijuana as a substitute for prescribing opioids. In research on overprescribing of opioids in states with medical and recreational marijuana laws, showed a 5.88% reduction in states with medical marijuana laws and a 6.38% decrease in states with adult use laws; including a decrease in Schedule II drugs of 7.79% and Schedule III to V by 10.40%.^12^ Their study found that Maryland did not have significant statistically discernable changes in opioid prescribing and Colorado had significantly lower prescribing rates.^12^ While in our research, Maryland did have a statistically different in the pre/post marijuana years, and Colorado still experienced a larger decrease in opioid distribution than Maryland or Utah.

Maryland actually had larger weights in every year, in every drug, when compared to Colorado and Utah. While Maryland did legalize medical marijuana, the law went into effect in June of 2014 with no real operations until late 2016. This rollout could have been less impactful due to bureaucratic and legal issues within the state leading to no significant changes in opioid prescribing.^12^ Maryland also had extremely high amounts of drugs used for OUD compared to both Utah and Colorado. There seems to be a higher rate of distribution of these medications, over the medications being used to treat pain.

Utah, while demographically similar to Colorado, had lower rates in every year and every drug than Colorado. Utah follows Colorado’s similar treads in Figure 1, 2, and 5. Utah’s drug distribution significantly differs from Colorado as represented in Figure 4. Colorado and Maryland showed a consistent increase in the distribution of drugs used for OUD. Utah has a consistent downward trend in OUD distribution. Utah follows a similar trend as Colorado and Maryland in distribution of pain medications. Overall, it appears that opioids used for both pain and OUD are not being distributed at an increased rate over the last decade. Complementing these findings, Utah has legalized medical use marijuana as of November 2018 with anticipated dispensary openings in January 2021. Continued research on these distribution treads will be needed.

Marijuana has also not been found to help in every respect in regard to opioid use. Olfson and colleagues found an association between illicit marijuana use with an increased incidence of nonmedical prescription opioid use (odds ratio = 5.8; 95% CI 4.23-7.90) using the National Epidemiological Survey on Alcohol and Related Conditions (NESARC).^23^ However, the NESARC findings may not apply to our study which evaluated an association between legalized marijuana and prescription opioids rather than between illicit marijuana use and nonmedical opioid use. Regardless, further research is needed to establish the relationship between marijuana and opioid use.

Marijuana may also be considered helpful in lowering the opioid overdose rate. In 2014, a study found a 24.8% reduction in deaths among states that had medical cannabis laws (MCLs)^8^ and fewer daily doses filled by patients (Medicare Part D population).^5^ Another report found a 6.5% reduction in overdose deaths after the legalization of recreational marijuana in Colorado.^9^ This represents an important reversal in the upward trend that Colorado was experiencing in opioid-related deaths. In combination of less opioid being distribution to the state, and concrete evidence that there has been a decrease in opioid-related deaths, Colorado’s legalized recreational marijuana shows an encouraging tread to treat people’s pain and keep them safe. Although, a more recent publication may have found a reversal in Bachhuber’s data over time. This study cites an association between state medical cannabis laws and opioid overdose reversed from −21% to +23% during the full timeframe of 1999-2017.^24^

Opioid prescribing practices were already beginning to decrease before any additional guidelines were released. When the CDC released the *Guideline for Prescribing Opioids for Chronic Pain* in March 2016, there was a substantial decrease in opioid prescribing practices.^25^ Research from this study confirms this finding. Colorado and Maryland experienced an overall decrease in opioid distribution, but Colorado’s decrease was larger. While the nation as a whole, was experiencing a decrease in opioid distribution, it was promising that Colorado’s greater decrease gives consideration to the potential impact of recreational marijuana.

## Limitations

There were some limitations with this study. There was a limited data range within ARCOS that was available. For enhanced data, it would have been advantageous to look at another decade. ACROS did not start posting data until 2000. Getting even closer to the beginning of the opioid epidemic may produce some great information about long term trends within the United States. This data also only looked at distribution data, not on a pharmacy or individual level. Improved data could come from tracking the drugs at the zip code level, but that is confounded by mail order pharmacies and internet pharmacies. Future pharmacoepidemiology research could also be adversely impacted if states elect for reimportation of prescription and controlled substances from Canada.

Confounding variables such as changing public opinion on marijuana use and related public health policies were not evaluated. During the timeframe of this study, neither Maryland nor Utah legalized recreational marijuana. However, Maryland passed a medical marijuana law in 2013 with sales of medical marijuana beginning in late 2017. Also, Maryland decriminalized possession of up to 10 grams of marijuana in 2014. Additionally, public opinion about marijuana use has been changing as evidenced in 2018 when Utah voters passed a medical cannabis law. This law which will become operational in Utah in 2021.^26^ These related marijuana policy changes could have had a confounding effect on medical opioid use.

It is unknown if patients actually reduce opioid use directly because of increased access to marijuana or if this association results from confounding by other factors. The intricacies of pain control and state level bureaucracies is tremendously multifactorial, and it makes it difficult to control every factor to isolate direct causation of marijuana on opioid distribution changes in an ecological investigation. The impact of guideline changes on the opioid distribution in 2016 may have confounded the results, not just the impact of marijuana within the three states. Further studies should focus more selectively on the subset of populations that are using recreational marijuana.

## Conclusion

In this study, we observed the differences in opioid distribution of eleven medications used for pain and OUD within Colorado, Utah, and Maryland from 2007 to 2017. Colorado, having legalized recreational marijuana, had a significant decrease in pain opioid distribution from 2007 to 2017. These findings further strengthen the need for additional research into the association of cannabis laws and prescription opioid use, since these findings do not show concrete evidence on the effects of marijuana. Combined with other novel research, marijuana may be useful for pain relief without dependency on opioids. There could be future utilization of guidelines to support decreasing prescribing practices for physicians and, we hope, the acceptance of marijuana as a treatment option for chronic pain. Health care providers and policymakers have the duty to consider other options for this epidemic, as it effects people across the country. There should also be a national level push for uniformity of marijuana policies under US federal law to at least allow for proper, concrete research on the overall health effects. If there is an initial reduction in opioid distributions in states with recreational marijuana laws, it is conceivable that opioid misuse, addiction, and overdose deaths could also fall. Therefore, it may be time to reconsider the practice of automatically discharging patients from pain treatment centers for positive marijuana screens, considering this use might actually reduce their overall opioid use. Further research should be considered on this topic and expanded to include more states and observe the long-term effects of recreational marijuana legalization.

## Acknowledgments

Data for this paper was obtained from the DEA as reported to ARCOS. The public availability of this data is valued. We could like to thank Iris Johnson at Geisinger for the article collection. No external funding was received for this research. BJP and SDN were supported by the Fahs-Beck Fund for Research and Experimentation.

---

**Table 1.** Table shows the relative comparison demographics between Maryland, Colorado, and Utah. Data was obtained from the US census 2017-2018.

