## Supplementary figures and images for "Prescription Opioid Distribution After the Legalization of Recreational Marijuana in Colorado, 2007-2017"

### Table 1

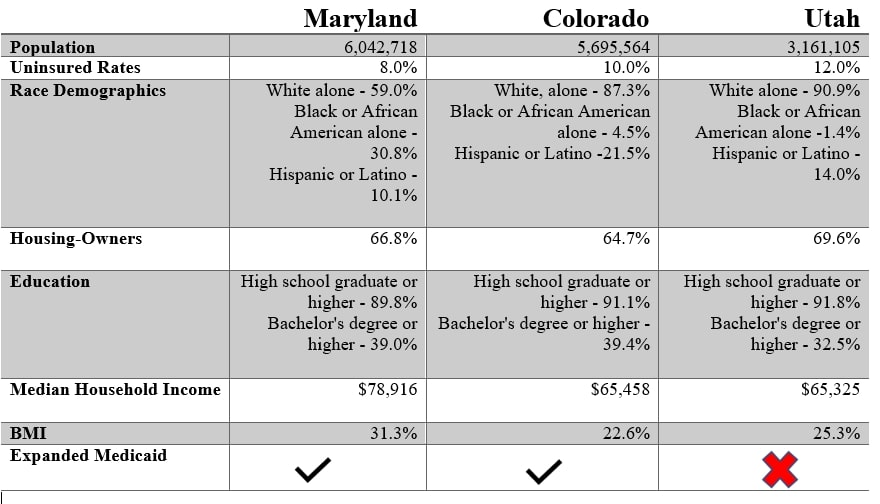
